# Host autophagosomes are diverted to a plant-pathogen interface

**DOI:** 10.1101/102996

**Authors:** Yasin F Dagdas, Pooja Pandey, Nattapong Sanguankiattichai, Yasin Tumtas, Khaoula Belhaj, Cian Duggan, Maria E Segretin, Sophien Kamoun, Tolga O Bozkurt

## Abstract

Filamentous plant pathogens and symbionts invade their host cells but remain enveloped by host-derived membranes. The mechanisms underlying the biogenesis and functions of these host-microbe interfaces are poorly understood. Recently, we showed that PexRD54, an effector from the Irish potato famine pathogen *Phytophthora infestans*, binds host protein ATG8CL to stimulate autophagosome formation and deplete the selective autophagy receptor Joka2 from ATG8CL complexes. Here, we show that during *P. infestans* infection, ATG8CL autophagosomes are diverted to the pathogen interface. Our findings are consistent with the view that the pathogen coopts host selective autophagy for its own benefit.

## Introduction

Plants intimately interact with a diverse range of pathogenic and symbiotic microbes, which typically produce specialized structures, such as haustoria and arbuscules, to invade the host cell space. Regardless of the interaction, these specialized structures are surrounded by membranes derived from the host endomembrane system (Koh et al., 2005; Le Fevre et al., 2015; Micali et al., 2011). These interfaces between the host and the invading microbes mediate inter-organismal communication enabling nutrient and macromolecule trafficking (Gutjahr and Parniske, 2013; Pumplin et al., 2012). However, our understanding of the origin and biogenesis of these host-microbe interfaces remains limited. In particular, the extent to which host accommodation membranes are shaped by the invading microbes is unclear.

Similar to many other filamentous plant pathogens, the potato late blight pathogen *P. infestans* produces haustoria, hyphal extensions that invaginate the host cell membrane (Whisson et al., 2016). Strikingly, the host accommodation membrane, also known as the extra haustorial membrane (EHM) contrasts sharply with the adjacent plasma membrane in both protein and lipid composition (Bozkurt et al., 2014; Lu et al., 2012; Schornack et al., 2009). Many of the tested membrane-localized proteins, notably cell surface immune receptors, are excluded from the EHM (Lu et al., 2012). A few exceptions include the membrane associated remorin protein REM1.3 and the vesicle fusion protein Synaptotagmin 1 (SYT1) (Bozkurt et al., 2014), which label distinct discrete membrane microdomains. Thus the EHM is not uniform, with multiple membrane sources probably contributing to its biogenesis (Bozkurt et al., 2015). The emerging model is that multiple trafficking pathways are diverted to haustoria with some degree of specificity. For example, whereas the vacuole-targeted late endocytic pathway marked by the Rab7 small GTPase RabG3C is diverted toward the EHM during *P. infestans* infection, another vacuole targeted protein Suc4 is not diverted (Bozkurt et al., 2015).

Autophagy is an evolutionary conserved membrane trafficking pathway that mediates removal or relocation of cytoplasmic components (Stolz et al., 2014). Autophagy involves formation of double membrane vesicles called autophagosomes coated with lipid bound ATG8 proteins. Autophagy cargoes are typically engulfed into autophagosomes and carried to the vacuole for cellular recycling (Lamb et al., 2013). We recently showed that a selective form of autophagy mediated by association of the plant autophagy cargo receptor Joka2 with the autophagy adaptor ATG8CL contributes to defense (Dagdas et al., 2016). To counteract this, *P. infestans* deploys an RXLR-WY type effector, PexRD54, carrying an ATG8 interacting motif (AIM) that mediates ATG8CL binding and depletes Joka2 from ATG8CL complexes (Dagdas et al., 2016). Interestingly, both PexRD54 and Joka2 preferably bind and stimulate formation of autophagosomes marked by potato ATG8CL over ATG8IL, highlighting the selective nature of the process (Dagdas et al., 2016). However, the fate of PexRD54 and Joka2 labeled autophagosomes during pathogen attack remains to be elucidated.

Here we investigated the subcellular dynamics of pathogen-targeted autophagy during *P. infestans* infection. We show that ATG8CL labeled autophagosomes accumulate around the haustoria, whereas ATG8IL labelled puncta rarely appear at the pathogen interface. Furthermore, an ATG8CL lipidation-deficient mutant remains cytoplasmic and fails to form perihaustorial puncta. Consistently, PexRD54 colocalizes with perihaustorial ATG8CL-autophagosomes in a manner that requires a functional AIM. Finally, Joka2 also accumulates at ATG8CL labeled perihaustorial puncta, indicating rerouting of autophagosomes towards the pathogen interface.

## Results and Discussion

### ATG8CL autophagosomes localize to haustoria in infected plants

To investigate subcellular dynamics of autophagy during infection, we first visualized transiently expressed GFP:ATG8CL in *N. benthamiana* epidermal cells during *P. infestans* infection. GFP:ATG8CL labeled autophagosomes frequently accumulated around the haustoria (73% of observations, *N*=60) labeled by the EHM marker REM1.3 (Figure 1A, Video 1-2, Figure 1 Supplementary Data). Unlike GFP:ATG8CL, autophagy deficient GFP:ATG8CLΔ mutant did not label any perihaustorial puncta (0%, *N*=30) and showed diffuse cytoplasmic signal similar to the GFP control (0% *N*=20) (Figure 1B-C, Figure 1 Supplementary Data). To test whether all autophagosomes are targeted toward the haustoria, we investigated subcellular localization of ATG8IL labeled autophagosomes. Unlike GFP:ATG8CL, GFP:ATG8IL labeled puncta rarely accumulated around the haustorium (20%, *N*=24) (Figure 1D, Figure 1 Supplementary Data). Taken together, these results demonstrate that during pathogen infection ATG8CL-autophagosomes are directed toward the pathogen interface and not the presumed vacuolar route.

**Figure 1.**
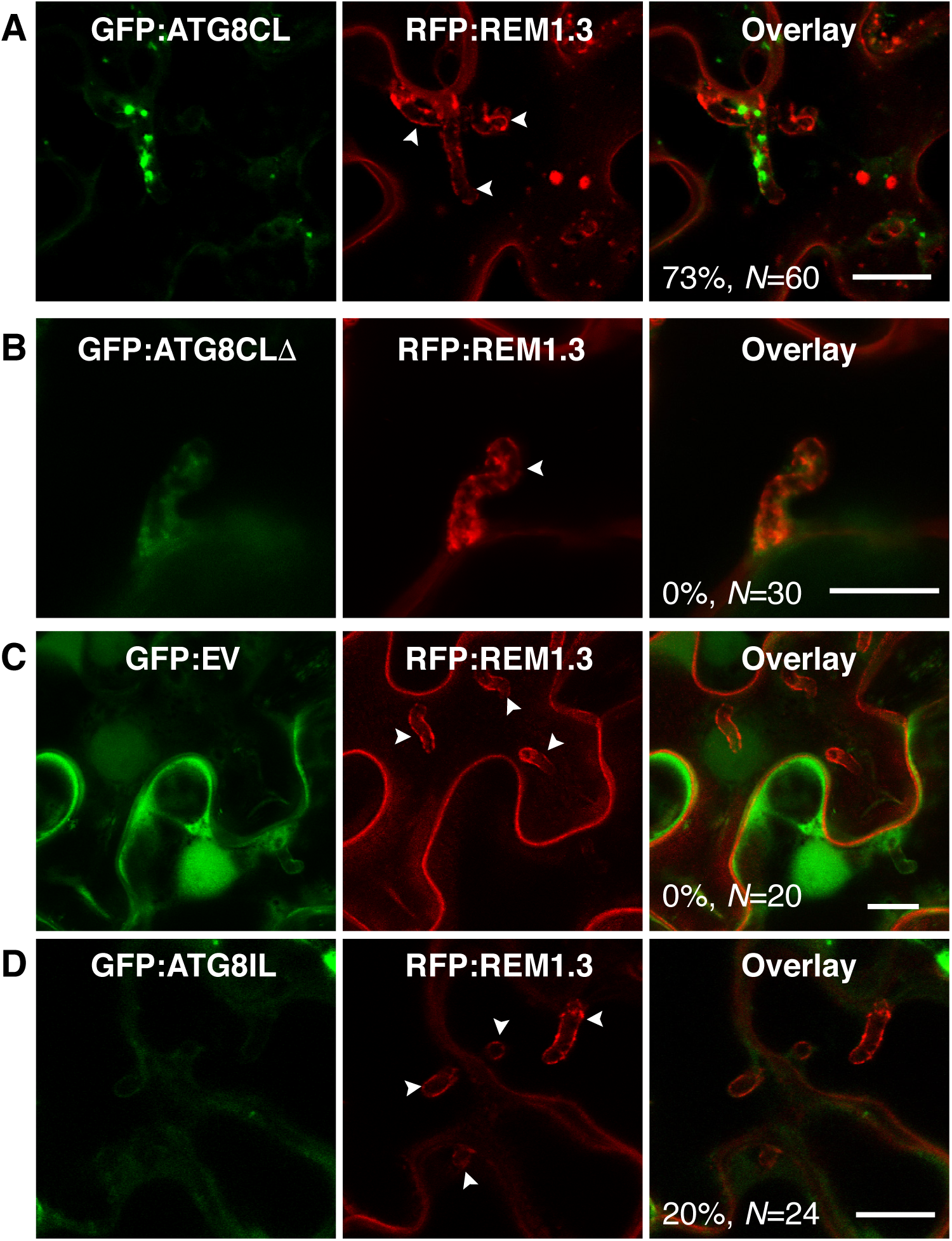
ATG8CL-autophagosomes accumulate around the haustoria. GFP:ATG8CL, GFP:ATG8CL, GFP:ATG8CLΔ, GFP:EV (empty vector) or GFP:ATG8IL are co-expressed with the EHM marker RFP:REM1.3 via agroinfiltration in *N. benthamiana* leaves infected with *P. infestans*. Confocal laser scanning microscopy was used to monitor the autophagosomes in haustoriated cells 3-4 days post infection (dpi). **(A)** GFP:ATG8CL frequently showed perihaustorial puncta whereas **(B)** autophagy deficient GFP:ATG8CLΔ mutant remained fully cytoplasmic similar to **(C)** GFP:EV. **(D)** GFP:ATG8IL, a divergent member of the ATG8 family, remained mostly cytoplasmic and rarely labeled perihaustorial puncta. Multiple optical sections that fully cover the haustoria are obtained to monitor perihaustorial puncta. Images shown are maximal projections of 15, 10 15, and 20 frames with 1μm steps for the top, upper middle, lower middle and bottom rows, respectively. Raw Z stacks are provided in Figure 1 Supplementary Data. Scale bars, 10 μm.

### PexRD54 accumulates at haustoria

Next, we set out to determine the subcellular localization of PexRD54 in infected cells. In haustoriated cells, GFP:PexRD54 frequently labeled perihaustorial puncta (70%, *N*=36) (Figure 2A, Figure 2 Supplementary Data). However, we rarely detected any puncta around the haustoria labeled by GFP:PexRD54^AIM2^ (5%, *N*=71), similar to the GFP control (0%, *N*=22) (Figure 2B-C, Figure 2 Supplementary Data). This suggested that ATG8CL binding is critical for recruitment of PexRD54 to the perihaustorial autophagosomes. To determine whether PexRD54 labeled vesicles are ATG8CL-autophagosomes, we co-expressed BFP:PexRD54 with GFP:ATG8CL in haustoriated *N. benthamiana* cells. We observed a full overlap between the two fluorescent signals, in contrast to BFP:PexRD54^AIM2^ and BFP:EV negative controls (100%, *N*=73 for PexRD54, 14%, *N*=35 for PexRD54^AIM2^ and 0%, *N*=29 for EV control) (Figure 2D-E-F, Figure 2 Supplementary Data). Although hardly observed (5%), detection of perihaustorial PexRD54^AIM2^ labeled autophagosomes suggests that this mutant can still weakly associate with ATG8CL *in vivo* or forms higher order molecular complexes with the host autophagy machinery. Overall, these findings demonstrate that the ATG8CL selective autophagy pathway that is targeted by *P. infestans* is diverted to the haustorial interface.

**Figure 2.**
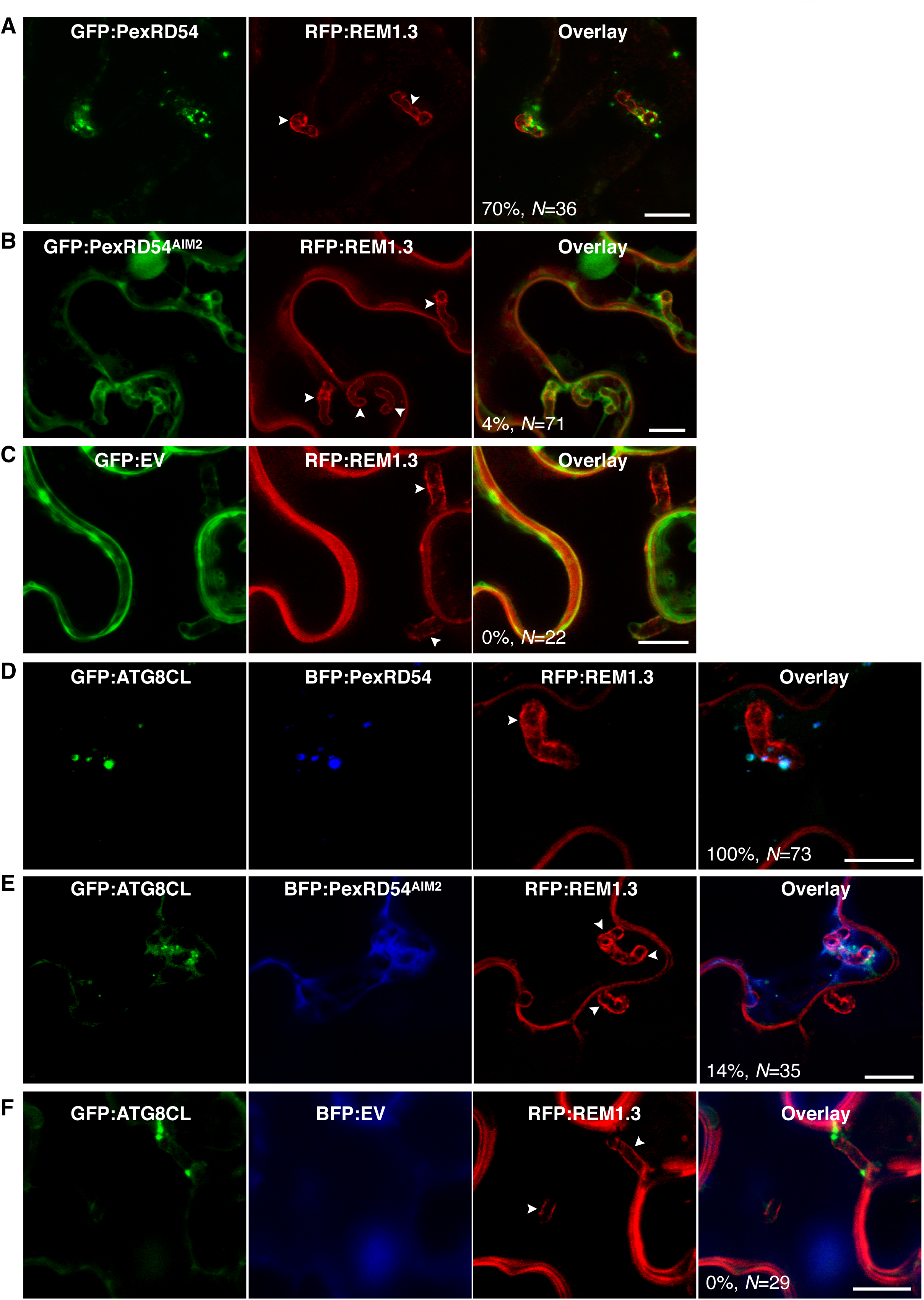
PexRD54 accumulates at perihaustorial ATG8CL-autophagosomes. (A-C) Confocal images of haustoriated *N. benthamiana* leaf epidermal cells marked by REM1.3 co-expressed with GFP:PexRD54, GFP:PexRD54^AIM2^ or GFP:EV control. GFP:PexRD54 showed frequent perihaustorial puncta (top panel) unlike its ATG8 interaction motif mutant PexRD54^AIM2^ (mid panel), which rarely labeled perihaustorial puncta and showed cytoplasmic distribution similar to GFP:EV (lower panel). Images shown are maximal projections of 11,15 and 25 frames with 1 μm steps from top to bottom rows, respectively. Arrowheads point to haustoria. Scale bars, 10 μm. Images were obtained 3-5 dpi. **(D-F)** Perihaustorial PexRD54 puncta overlaps with ATG8CL-autophagosomes. In haustoriated cells marked by RFP:REM1.3, GFP:ATG8CL labeled autophagosomes fully overlapped with perihaustorial BFP:PexRD54 puncta (top panel). In contrast, BFP:PexRD54^AIM2^ mainly remained cytoplasmic and mostly did not show perihaustorial puncta that overlap with ATG8CL autophagosomes (mid panel), similar to BFP:EV control (lower panel). Images shown are maximal projections of 10, 5, 8, frames with 1μm steps from top to bottom rows, respectively. Raw Z stacks are provided in Figure 2 Supplementary Data. Arrowheads point to haustoria. Scale bars, 10 μm. Images are obtained 3-4 dpi.

### Host autophagy cargo receptor Joka2 accumulates at haustoria

We also investigated subcellular trafficking of Joka2-autophagosomes in haustoriated cells. Joka2:BFP labeled autophagosomes frequently accumulated around the haustoria (92%, *N*=50) (Figure 3A, Figure 3 Supplementary Data) unlike BFP vector control (0%, *N*=37) (Figure 3C, Figure 3 Supplementary Data). Intriguingly, Joka2^AIM^:BFP mutant also labeled perihaustorial puncta, although its frequency was lower (74%, *N*=42) compared to Joka2:BFP (Figure 3B, Figure 3 Supplementary Data). To validate that Joka2 labeled compartments are indeed autophagosomes, we co-expressed BFP:Joka2, BFP:Joka2^AIM^ or BFP:EV with GFP:ATG8CL in haustoriated *N. benthamiana* cells marked by RFP:REM1.3. BFP:Joka2 fluorescent signal fully overlapped with GFP:ATG8CL labeled perihaustorial autophagosomes (100%, *N*=140) unlike the BFP:EV (0%, *N*=20) indicating Joka2 localizes to perihaustorial ATG8CL autophagosomes (Figure 3D-F, Figure 3 Supplementary Data). Surprisingly, Joka2:BFP also labeled puncta that did not show any GFP:ATG8CL fluorescence suggesting that Joka2 also labels compartments that are not ATG8CL-autophagosomes (Figure 3B, Figure 3 Supplementary Data). Consistent with this, Joka2^AIM^:BFP produced fluorescence signal at discrete puncta that rarely coincided with perihaustorial ATG8CL-autophagosomes (19%, *N*=37) (Figure 3E, Figure 3 Supplementary Data). Like mammalian autophagy cargo receptors, Joka2 forms oligomers, and this most likely accounts for recruitment of Joka2^AIM^:BFP to ATG8CL-autophagosomes. Nevertheless, this shows that Joka2 labels at least two perihaustorial compartments; one including ATG8CL-autophagosomes that requires a functional AIM; and another in which the AIM is dispensable. These results further illustrate that ATG8CL/Joka2 mediated antimicrobial autophagy is targeted to the haustorial interface.

**Figure 3.**
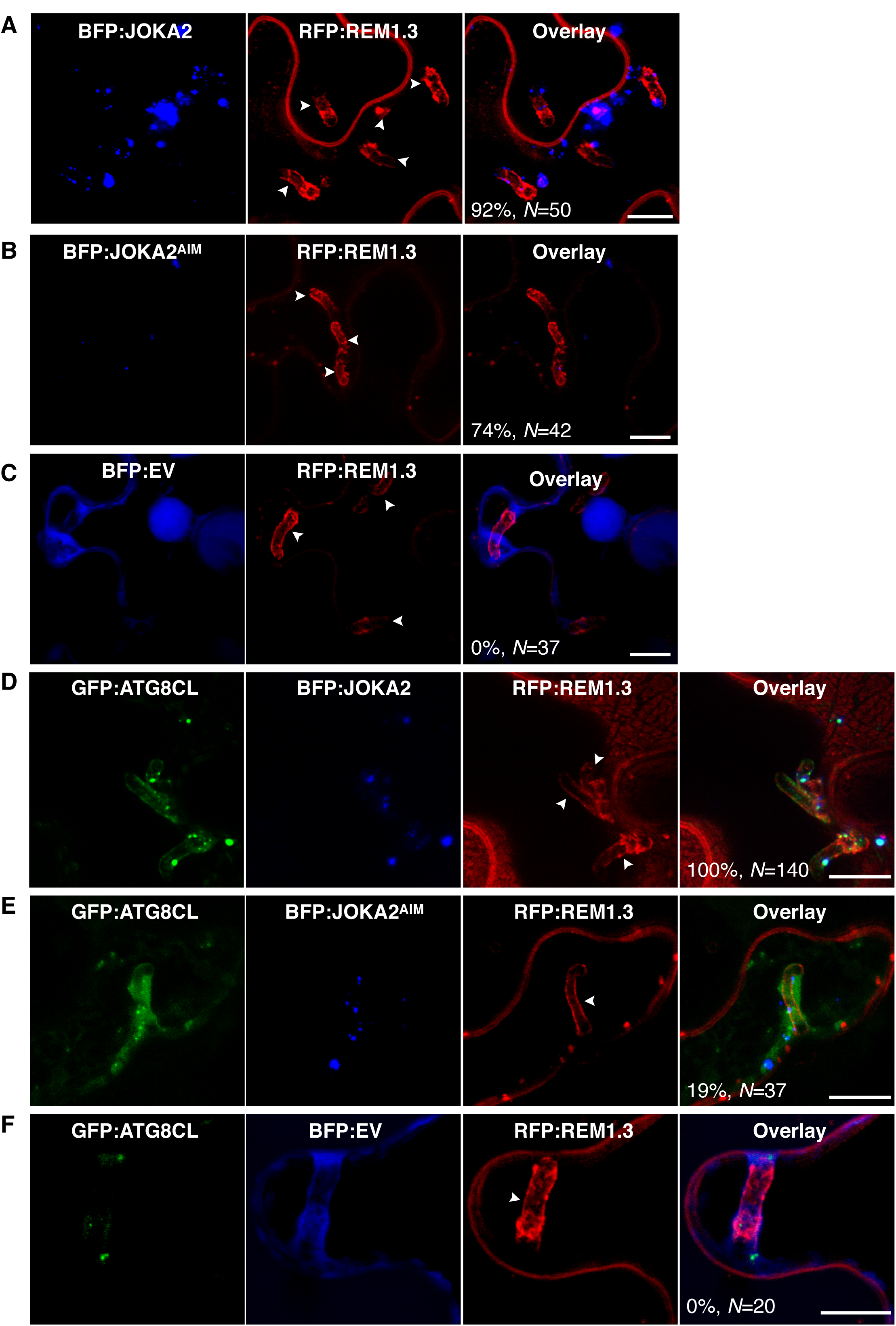
Joka2-mediated autophagy is directed toward the haustoria. (A-C) Confocal microscopy of *P. infestans* infected *N. benthamiana* leaf epidermal cells expressing Joka2:BFP, Joka2^AIM^:BFP mutant or BFP:EV control. Both Joka2:BFP (top panel) and Joka2^AIM^:BFP (mid panel) displayed perihaustorial puncta although the frequency of the later was much lower. Consistently, no punctate structures were labeled by the BFP:EV control (lower panel) around the haustoria. Images shown are maximal projections of 8,14 and 7 frames with 1 μm steps from top to bottom rows, respectively. Arrowheads point to haustoria. Scale bars, 10 μm. Images were obtained 3-4 dpi. **(D-F)** Joka2 localizes to ATG8CL-autophagosomes around the haustoria. GFP:ATG8CL labeled perihaustorial autophagosomes fully overlapped with the Joka2:BFP labeled puncta, whereas Joka2^AIM^:BFP generally did not label GFP:ATG8CL autophagosomes similar to BFP:EV control. Images shown are maximal projections of 10, 10 and 8, frames with 1μm steps from top to bottom panels, respectively. Raw Z stacks are provided in Figure 3 Supplementary Data. Arrowheads point to haustoria. Scale bars, 10 μm. Images are obtained 3 dpi.

### Host autophagosomes are diverted to a plant-pathogen interface

In this study, we employed high-resolution microscopy to monitor the course of defense related autophagy in *N. benthamiana* cells during *P. infestans* infection. We show that autophagosomes labeled by the core autophagy protein ATG8CL, as well as the pathogen secreted effector protein PexRD54 and the host autophagy cargo receptor Joka2 are transported from the cytoplasm to the EHM. This reveals a novel trafficking route from the cytoplasm to the pathogen interface and expands our understanding of the biogenesis of the EHM (Bozkurt et al., 2015; Bozkurt et al., 2014; Lu et al., 2012).

We propose a model in which pathogen-manipulated autophagosomes are channeled toward the haustorial interface. A provocative hypothesis is that the PexRD54-ATG8C autophagosomes carry a distinct cargo that substitutes the defense related cargo to redirect molecules towards the pathogen. Such pathogen modified double-layered autophagosomes could fuse with the EHM discharging single layered vesicles into the extrahaustorial matrix. Fusion of autophagosomes with the EHM could provide a membrane source for EHM biogenesis and may account for the extracellular vesicles observed in various host-microbe interfaces (Deeks and Sánchez-Rodríguez, 2016). Further studies are required to determine the precise mechanisms that govern autophagosome transport to the haustorial interface and its impact in pathogenicity. Moreover, identifying the nature of the autophagosome cargo sequestered by PexRD54 and Joka2 should help better understanding the role of selective autophagy in this host-microbe interaction.

## Materials and Methods

### Molecular cloning and plasmid constructs

GFP:ATG8CL, GFP:ATG8CLΔ, GFP:ATG8IL, GFP:PexRD54, GFP:PexRD54^AIM^, GFP:EV and RFP:REM1.3 constructs were previously described by Dagdas et al. (2016). All other blue fluorescent protein (BFP) fusion constructs were generated in this study. The vector for N-terminal BFP fusion was derived from pK7WGF2 plasmid (Karimi et al., 2002) by excising a fragment from the backbone with EcoRV digestion then replacing it with a custom synthesized fragment containing tagBFP sequence followed by linker sequence (GGATCTGCTGGATCTGCTGCTGGATCTGGAGAATTT) and EcoRV restriction site (where the gene of interest will be inserted) (Eurofins Genomics). Similarly, the vector for C-terminal BFP fusion was also derived from pK7WGF2 plasmid but by inserting PCR fragments containing EcoRV restriction site (where the gene of interest will be inserted) followed by linker sequence (GGATCTGCTGGATCTGCTGCTGGATCTGGAGAATTTGGATCA) and tagBFP sequence amplified from N-terminal BFP fusion vector using primer pairs GA_35s_F with Cterm_BFP_Prom_R and Cterm_BFP_F with Cterm_BFP_R. Then, BFP:PexRD54, BFP:PexRD54^AIM2^, Joka2:BFP and Joka2^AIM^:BFP constructs were generated by Gibson assembly of each gene PCR fragment into EcoRV digested BFP vectors (N-terminal fusion for PexRD54 and PexRD54^AIM^, C-terminal fusion for Joka2 and Joka2^AIM^). All genes were amplified from existing constructs previously described by Dagdas et al. (2016) using primer pairs GA_RD54_F with GA_RD54_R for PexRD54, GA_RD54_F with GA_LIR2_R for PexRD54^AIM^ and GA_Joka2_BFP_F with GA_Joka2_BFP_R for both Joka2 and Joka2^AIM^. All primers used in this study are listed in Supplemental Table 1.

### Confocal Microscopy

Imaging was performed using Leica SP5 resonant inverted confocal microscope (Leica Microsystems) using 63x water immersion objective. All microscopy analyses were carried out on live leaf tissue 3-4 days after agroinfiltration. Leaf discs of *N. benthamiana* were cut and mounted onto Carolina observation gel (Carolina Biological Supply Company) to minimize the damage. Specific excitation wavelengths and filters for emission spectra were set according to Lu et al. (2012). BFP, GFP and RFP probes were excited using 405, 488 and 561 nm laser diodes and their fluorescent emissions detected at 450-480, 495–550 and 570–620 nm, respectively. Sequential scanning between lines was done to avoid spectral mixing from different fluorophores and images acquired using multichannel. Maximum intensity projections of Z-stack images were presented in each figure. Z-stack sections were processed to enhance image clarity, sections that caused blurriness (top and bottom ones), were removed for generation of maximum intensity projections. Representative raw Z-stacks are provided in Supplementary data files 1,2 and,3. Image analysis was performed using ImageJ (1.50g) and Adobe Photoshop (CS6).

### Transient gene-expression assays in *N. benthamiana*

Transient gene-expression was performed *in planta* by infiltration of leaves of 3-4 week old *N. benthamiana* with cultures of *Agrobacterium tumefaciens* GV3101 strain carrying T-DNA constructs, as previously described (Bozkurt et al., 2011). Transient co-expression assays were carried out by mixing equal ratios of *A. tumefaciens* carrying the plant expression constructs in agroinfiltration medium [10 mM MgCl_2_, 5 mM 2-(N-morpholine)-ethanesulfonic acid (MES), pH 5.6] to achieve a final OD_600_ of 0.2.

### Biological Material

*N. benthamiana* plants were grown and maintained in a greenhouse with high light intensity (16 hours light/ 8 hours dark photoperiod) at 22-24°C. *P. infestans* strain 88069 cultures (van West et al., 1998) were grown and maintained on rye sucrose agar medium at 18°C in the dark for 12-14 days, as described elsewhere (Song et al., 2009) prior to use for infection of *N. benthamiana.* Zoospores were released from sporangia by addition of cold water and incubation at 4°C for 1-2 hours adjusting dilution to 50,000 spores/ml. Infection of agroinfiltrated leaves was carried out by addition of 10 μl droplets containing zoospores as described previously (Saunders et al., 2012; Song et al., 2009) with the exception that infection was carried out on live plants, incubating inoculated plants in humid growth chambers.

## Acknowledgements

We thank members of the Bozkurt and Kamoun Labs for helpful suggestions. This project was funded by the Gatsby Charitable Foundation, European Research Council (ERC), and Biotechnology Biological Sciences Research Council (BBSRC).

## Author Contributions

YFD, Conception and design, Acquisition of data, Analysis and interpretation of data, Drafting or revising the article

PP, Conception and design, Acquisition of data, Analysis and interpretation of data, Drafting or revising the article

NS, Conception and design, Acquisition of data, Analysis and interpretation of data, Drafting or revising the article

YT, Conception and design, Acquisition of data, Analysis and interpretation of data,

CD, Acquisition of data, Analysis and interpretation of data, Drafting or revising the article MES, Acquisition of data, Analysis and interpretation of data

SK, Conception and design, Analysis and interpretation of data, Drafting or revising the article

TOB, Conception and design, Acquisition of data, Analysis and interpretation of data, Drafting or revising the article

## Video 1-2. ATG8CL-autophagosomes accumulate around the haustoria

GFP:ATG8CL is co-expressed with the EHM marker RFP:REM1.3 via agroinfiltration in *N. benthamiana* leaves infected with *P. infestans*. Confocal laser scanning microscopy was used to monitor the autophagosomes in haustoriated cells 3-4 dpi. In Video 1, RFP channel represents maximum projection of Z-stack of images illustrated in Video 2 to better illustrate the haustorium.

